# Atlas-scale spatially aware clustering with support for 3D and multimodal data using SpatialLeiden

**DOI:** 10.64898/2026.02.27.708246

**Authors:** Niklas Müller-Bötticher, Alexander Malt, Paul Kiessling, Roland Eils, Christoph Kuppe, Naveed Ishaque

**Author notes:** Corresponding author **Correspondence**, Correspondence should be sent to. Equal contribution (Niklas Müller-Bötticher, Alexander Malt).

## Abstract

Here we extend SpatialLeiden, our spatial clustering algorithm, to enable generalised atlas-scale multi-sample, 3D serial-section, and multimodal spatial omics via flexible neighbour-graph multiplexing on batch-corrected latent spaces. It delivers coherent domains aligning with brain atlases across >100 samples, stable 3D reconstruction of cancer tissue structures, and integrated multimodal features, outperforming specialized tools in modularity and scalability on standard hardware. SpatialLeiden is compatible with scverse for broad and intuitive adoption.

## Introduction

Spatially resolved transcriptomics (SRT) measures gene expression in intact tissue context, yielding high-dimensional datasets that reveal cellular organization but can be difficult to interpret.^1^ Clustering by transcriptional similarity and spatial proximity is essential for spatial atlas construction, tissue architecture analysis, and marker discovery.^2^ A wide range of clustering algorithms has been developed for spatial transcriptomics data, with approaches ranging from graph convolutional networks to Markov random fields. Spatial Leiden is comparatively simple: it integrates spatial information via neighbourhood graphs alongside the standard gene expression *k*-NN graph, rendering the well-established Leiden community detection algorithm spatially aware.^3^

Large-scale multi-sample, 3D volumetric, and multimodal spatial omics studies boost statistical power and biological insight. Batch correction methods from single-cell RNA-seq (scRNA-seq) learn shared latent spaces to align biological states while minimising technical variation.^4^ Serial sectioning enables volumetric SRT with 3D (*x,y,z*) neighbourhoods that capture inter-slice continuity.^5^ Spatial multi-omics platforms jointly profile transcripts, proteins, chromatin, immune repertoire, and imaging features to reveal complex cellular states beyond transcriptomics alone.^6–17^ Yet this complexity introduces cross-slice variability, complicating spatial domain alignment and yielding inconsistent results.^18^

Here, we extend SpatialLeiden with multimodal support and workflows for multi-sample and serially sectioned data. This enables robust, efficient clustering of batch-corrected latent spaces and multi-omic layers for integrative spatial clustering across atlas-scale datasets or 3D volumes while preserving biological and spatial structure.

## Results

### Flexible multi-sample spatially aware clustering

SpatialLeiden applies Leiden community detection to a collection of neighbourhood graphs. It is agnostic to the creation of this graph and previous processing steps, which allows the user to freely select the optimal integration strategy (**Supplementary Figure 1a**). We demonstrate this flexibility on the dorsolateral prefrontal cortex (DLPFC) Visium dataset by integrating latent spaces with scVI,^19^ Harmony,^20^ or BBKNN.^21^ For within-patient and cross-patient integration scVI maintains clustering accuracy as indicated by the adjusted Rand index and normalized mutual information across samples while also enabling joint clusters (**Supplementary Figure 1**).

### SpatialLeiden processes atlas-scale datasets

SpatialLeiden scales to atlas-level datasets, as demonstrated by clustering all ∼150 serial MERFISH sections from the Allen Brain Cell (ABC) Atlas.^22^ Identified clusters align closely with the Common Coordinate Framework (CCF) parcellations^23^ (**Figure 1a**). Increasing resolution further refines known multi-scale spatial structures (**Figure 1b**) like isocortical layer separation (**Supplementary Figure 2**). The clusters are consistent with expected cell type enrichment and CCF parcellation indices, showcasing the biological validity of the approach (**Figure 1 c,d**). Compared to non-integrated clustering (**Supplementary Figure 3**), integrated clusters exhibit smooth composition transitions across brain slices (**Figure 1e**). SpatialLeiden is runtime- and memory-efficient, allowing processing of atlas-scale datasets even on high-end notebooks (**Figure 1f**).

**Figure 1:**
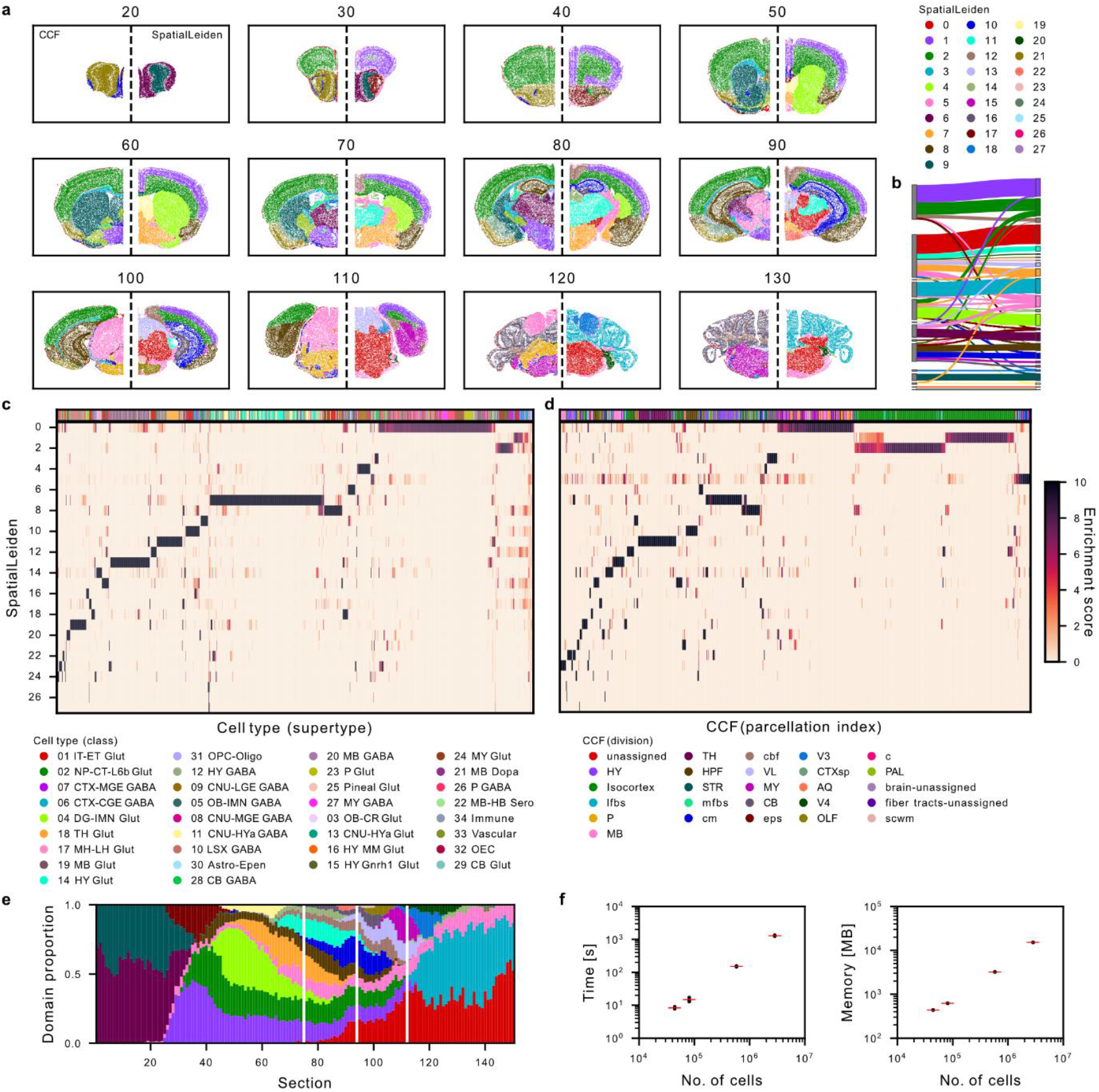
SpatialLeiden identifies domains in whole brain cell atlas. **a**, CCF annotation and SpatialLeiden domains for representative sections. **b**, Movement of cells between domains for resolution 0.5 (left) and resolution 1 (right). **c–d**, Cell type classes (**c**) and CCF structures (**d**) are enriched within distinct SpatialLeiden domains. **e**, The SpatialLeiden domains are spatially continuous across the serial sections. **f**, Runtime and memory for clustering subsampled sections of the dataset. Red line indicates the mean of three runs. CCF, common coordinate framework.

### 3D integration of stitched serial sections

SpatialLeiden can incorporate *z*-axis information directly into its spatial neighbourhood graph, enabling 3D clustering of stitched serial sections or segmented data of thick sections. Applied to aligned Open-ST metastatic lymph node data^24^ it revealed 3D cellular neighbourhoods spanning across sections (**Figure 2a, Supplementary Figure 4**). The domain composition remains stable across profiled sections (**Figure 2c,d**), with distinct cell type profiles per domain highlighting tumour microenvironment organization (**Figure 2b**).

**Figure 2:**
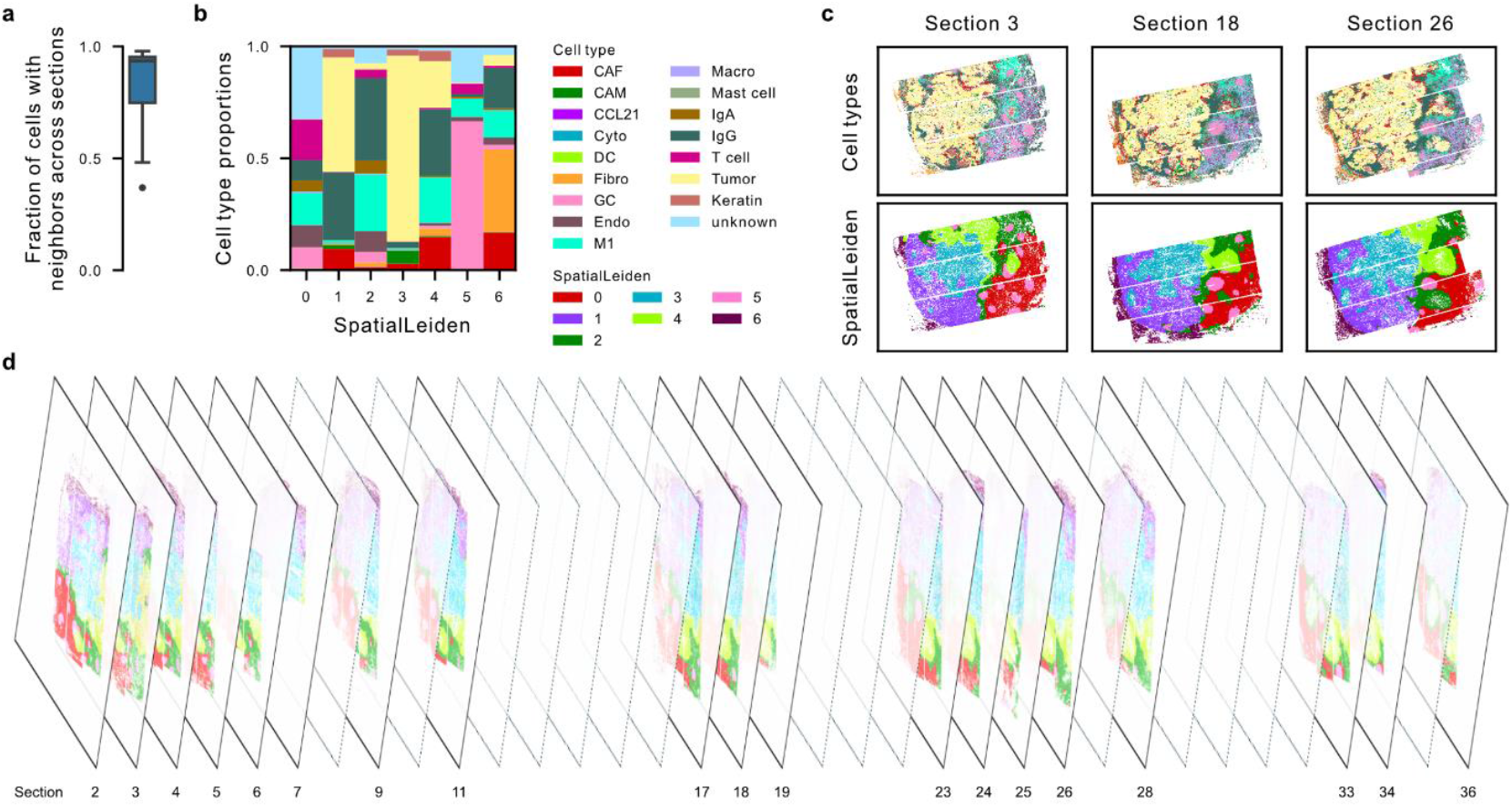
3D SpatialLeiden using spatially aligned serial sections. **a**, Fraction of cells with at least one spatial neighbour in a different section. **b**, Cell type composition of the spatial domains. **c**, Example sections with cells coloured by cell type (top) and spatial domain (bottom). **d**, Spatial domains across serial, 3D-stacked sections (only indicated sections were processed with Open-ST). Box plots show the median and the 25^th^–75^th^ percentiles, with whiskers extending to the most extreme data points within 1.5× the interquartile range. CAF, cancer-associated fibroblast; CAM, cancer-associated macrophage; CCL21, CCL21^+^ endothelial cells; Cyto, cytotoxic T cell; DC, dendritic cell; Endo, endothelial venule; Fibro, fibroblast; GC, germinal center IgM plasma cell; IgA, IgA plasma cell; IgG, IgG plasma cell; M1, M1 macrophage; Macro, macrophage; Keratin, keratin pearl.

### Multimodal data integration

SpatialLeiden’s multimodal extension treats each omic or imaging layer as an extendable adjacency element within a MuData structure,^25^ enabling joint clustering via customisable layer weights. Unlike methods limited by modality count, SpatialLeiden flexibly scales to multiple layers (**Figure 3a**). Applied to a synthetic dataset,^1^ SpatialLeiden showed excellent performance (**Supplementary Figure 5**). SpatialLeiden’s compatibility with scverse enables easy integration with LazySlide^26^ for handling and processing of imaging modalities. In a Visium HD colorectal cancer (CRC) dataset with matched histology,^27^ individual modalities delineated particular features, where histology refined the smooth muscle clusters (**Supplementary Figure 6**), while SRT separated dysplasia from neoplasia (**Supplementary Figure 7**). Joint SpatialLeiden clustering of both modalities simultaneously resolved smooth muscle refinement and dysplasia-neoplasia separation (**Figure 3b–d**).

**Figure 3:**
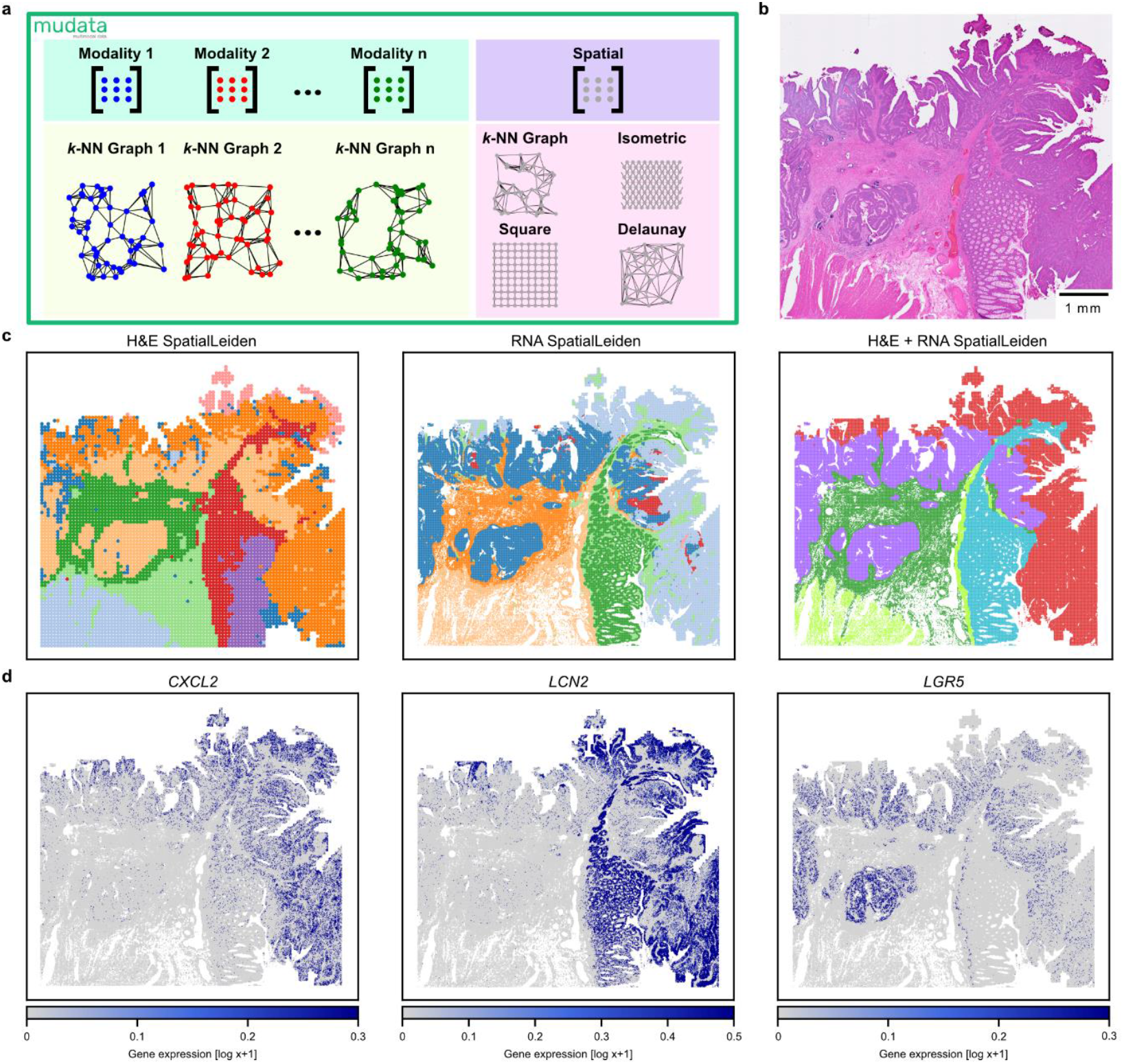
Multimodal SpatialLeiden. **a**, Multimodal data and associated k-NN graphs are integrated in a MuData object, which serves as input for multimodal SpatialLeiden. **b**, H&E section from CRC Visium HD slide. **c**, SpatialLeiden clustering on tiled H&E (left), binned RNA (middle), and multimodal integration with a shared bin size (right). **d**, Normalized gene expression of CXCL2, LCN2, and LGR5 highlight dysplastic (red) and neoplastic (purple) regions identified exclusively from RNA data.

## Discussion

SpatialLeiden demonstrates several important advantages. It seamlessly scales to any number of modalities with matched cells/spots as adjacency components, offering greater extensibility compared to existing methods such as SpatialGlue.^1^ This extensibility enables researchers to tailor the framework to diverse experimental setups and data types, broadening its applicability across spatial biology studies. Furthermore, it naturally extends to handle multi-slice data and 3D serial sections, addressing growing demand for volumetric spatial analysis. By focussing sole on neighbour graphs and clustering, the approach remains preprocessing agnostic, allowing users to leverage custom preprocessing steps such as future spatially-aware batch correction.

We have previously shown that existing spatial clustering methods fail to scale to large datasets.^28^ SpatialLeiden demonstrates linear scaling, maintaining computational feasibility when analysing millions of cells and hundreds of samples. This becomes increasingly critical as spatial omics and imaging datasets grow exponentially.

Benchmarking current multimodal and 3D spatial omics datasets remains challenging. We found simulated data^1,29^ to be overly simplistic, and multimodal data from real tissues remains scarce, of low quality and/or limited number of samples. With an increasing number of modalities, manual weighting of modality relevance introduces additional complexity in multimodal integration. Furthermore, 3D clustering accuracy relies heavily on precise registration of serial sections. Misalignment or registration artifacts may distort spatial relationships, leading to erroneous cluster assignments and misleading biological interpretations.

## Methods

Data was analysed using Python (v3.12.12) with the following package versions (unless otherwise specified): anndata v0.10.9, geopandas v1.1.2, leidenalg v0.10.2, mudata v0.3.3, pandas v2.1.1, scanpy v1.10.4, scikit-learn v1.8.0, scipy v1.14.1, snapatac2 v2.8.0, spatialleiden v0.4.0, squidpy v1.5.0, torch v2.10.0. All relevant function calls were seeded with a random state for reproducibility. Unspecified parameters are used with default values.

### Spatially-aware Leiden multiplex (SpatialLeiden)

For multi-sample and 3D clustering all samples were concatenated in a single AnnData^30^ object. Batch integration was performed prior to building the *k-*NN graph in the gene expression space. For 3D clustering, the physical neighbour graph is build using the 3D coordinates. This requires the serial sections to be spatially aligned before graph construction.

For multimodal SpatialLeiden we use MuData^25^ objects to ensure that the observations (and therefore nodes of each graph) are matched. Each modality is stored as an AnnData object and the *k-*NN graph of each modality’s feature space can be generated using scanpy^31^ as described below. The physical neighbour graph can be generated using squidpy^32^ (same as for the single-modality case). For clustering the parameters (including resolution, directedness of the graph, and the weights for each modality) can be set for each modality separately or jointly for all.

### Preprocessing

The data was preprocessed using scanpy by filtering genes that are expressed in at least 10 cells (scanpy.pp.filter_genes) and selecting 2,000 highly variable genes using batch information if available (scanpy.pp.highly_variable_genes with flavor ‘seurat_v3’). Subsequently the data was normalized (scanpy.pp.normalize_total), log-transformed (scanpy.pp.log1p), scaled without zero-centering unless specified (scanpy.pp.scale), and the first 30 principal components calculated (scanpy.pp.pca).

### Batch integration

BBKNN^21^ (bbknn v1.6.0) was run using scanpy.experimental.pp.bbknn using the sample information as batch variable. To integrate data with Harmony^20^ (harmonypy v0.2.0), sample (and patient) information was used to correct the PCA transformed gene expression. Batch correction with scVI^19^ (scvi-tools v1.2.2.post2) was performed on the raw counts using sample information as batch variable (and patient information if applicable as categorical covariates). A negative binomial distribution was used as gene likelihood with 2 hidden layers for the encoder and decoder networks.

### Neighbourhood graph generation

The 15 nearest neighbours in the (batch-corrected) gene expression latent space were identified to obtain the undirected *k-*NN graph (scanpy.pp.neighbors). The physical neighbours graph was calculated with squidpy.gr.spatial_neighbors using (i) 10 neighbours with coord_type ‘generic’ for the MERFISH ABC Atlas and Open-ST lymph node (ii) 6 neighbours with coord_type ‘grid’ for the Visium DLPFC dataset and (iii) 8 neighbours with coord_type ‘grid’ for the Visium HD CRC dataset and the simulated data. In case of Open-ST and MERFISH datasets the connectivities of the spatial neighbours graph were calculated using spatialleiden.distance2connectivity (grid-aligned neighbour structures are treated as equidistant and therefore have the same connectivity). The resulting graphs are undirected in the case of regular grids and directed in the case of generic coordinate systems.

### Visium dorsolateral prefrontal cortex

The DLPFC samples were processed as described above. The data was zero-centred and scaled. For BBKNN 5 neighbours were used in the case of patient-wise integration of the and 3 neighbours when integrating across patients to avoid the *k*-NN graph to have too many edges compared to the other batch integration methods.

SpatialLeiden was run using the spatialleiden.search_resolution function to obtain the same number of clusters as defined in the ground truth annotation. The layer ratio was set to 0.8 in the single sample case and for BBKNN, and to 1.2 otherwise (to counteract the effect that more neighbours of the gene expression *k*-NN graph will be across sections and achieve comparable spatial continuity of the identified domains).

### MERFISH Allen Brain Cell Atlas

The sections of the ABC Atlas were integrated with scVI and clustered with spatialleiden using a layer ratio of 2.5 for 5 iterations. The resolution was set to 1 unless otherwise specified. The enrichment score was calculated as fold change of the cell density of an annotation (cell type or CCF parcellation index) within a spatial domain to the density across all domains following the approach in Zhang *et al*.^33^

### Open-ST metastatic lymph node

Open-ST metastatic lymph node data was integrated using scVI. To avoid long distance neighbours in the spatial graph, all connections > 300 (∼100 µm) were removed before calculating the connectivities with distance2connectivity. Clustering with spatialleiden was performed with a layer ratio of 2 and a resolution of 0.7 for 10 iterations.

### Visium HD colorectal cancer

The H&E-stained whole slide image (WSI) was transformed to an image pyramid using pyvips (v3.0.0 with libvips v8.17.2) and processed using LazySlide^26^ (v0.9.5 with wsidata v0.8.1) to identify tissue regions (lazyslide.pp.find_tissues at level 3) without hole detection to preserve all tissue regions. To focus the analysis on the area overlapping with SRT data, the tissue boundary was cropped to a predefined region of interest (40550–65300×0–22750 pixels). The resulting tissue region was partitioned into 256×256 pixel tiles (∼69 µm, lazyslide.pp.tile_tissues). Feature vectors were then extracted for each tile using the UNI2 model^34^ (lazyslide.tl.feature_extraction).

Tile-level feature matrices were scaled without zero-centering (scanpy.pp.scale) and the first 15 principal components calculated (scanpy.pp.pca). The 15 nearest neighbours in the latent space were identified to obtain the *k-*NN graph (scanpy.pp.neighbors).

Pre-defined resolutions were used to perform tile-level clustering (scanpy.tl.leiden). Spatially informed clustering was performed using spatialleiden with a layer ratio of 0.5 and a resolution of 0.4.

The 8 µm SRT bins were preprocessed as described in the Preprocessing chapter. The bin centroids were computed and mapped to tiles (geopandas.sjoin) to generate a mapping linking each bin to its corresponding tile.

Neighbourhood connectivity tile graphs were recalculated with 4 neighbours and a gaussian kernel (scanpy.pp.neighbors), the tiles were mapped to the bins, and the resulting connectivity matrices were downsampled to a maximum of 15 neighbours per row. The binned RNA data and corresponding tile features were merged into a MuData object.

Multimodal clustering was performed (spatialleiden_multimodal) with a resolution of 0.55 incorporating RNA, H&E, and spatial layers, with layer weights of 1.5, 0.5, and 2, respectively.

### ARI and NMI calculation

The adjusted Rand index and normalized mutual information were calculated using the adjusted_rand_score and normalized_mutual_info_score functions in scikit-learn^35^ with all parameters at default values.

### Scalability benchmark

To evaluate the runtime and memory of the clustering with spatialleiden (v0.3.0), the number of sections used from the ABC Atlas was randomly downsampled and the gene expression neighbours graph recalculated. SpatialLeiden was run for 3 iterations. GNU time with elapsed time and memory (maximum resident set size) was used to measure runtime characteristics.

## Declarations

### Data Availability

All data used in this publication is publicly available; DLPFC profiled using the 10x Genomics Visium platform^36^ (https://research.libd.org/spatialLIBD/), Allen Brain Cell Atlas profiled using MERFISH^22^ (manifest 20250131 downloaded with abc_atlas_access v3.1.3, https://github.com/alleninstitute/abc_atlas_access), metastatic lymph node sequenced with Open-ST^24^ (GEO accession no. GSE251926), simulated data from the SpatialGlue publication^1^ (https://zenodo.org/records/10362607), and the Visium HD CRC dataset^27^ from 10x Genomics website (https://www.10xgenomics.com/datasets/visium-hd-cytassist-gene-expression-libraries-of-human-crc).

### Code Availability

The code to reproduce all results from this study is publicly available on GitHub; https://github.com/HiDiHlabs/SpatialLeiden2-Study. The SpatialLeiden Python package is available on GitHub, PyPI, and Bioconda as free and open-source software licensed under an GPLv3 license; https://github.com/HiDiHlabs/SpatialLeiden.

### Author contributions

N.I., N.M.B. conceived and designed the study. N.M.B., A.M., P.K. implemented code. A.M. performed code review. N.M.B., A.M. performed data analysis. N.I., N.M.B., A.M. interpreted and analysed results. N.I., N.M.B., A.M. wrote the manuscript. All authors proofread and corrected the manuscript. All authors contributed to the article and approved the submitted version.

## Acknowledgements

This project was conceptualized during the de.NBI BioHackathon SpaceHack 3.0 in Berlin (December 2024), we thank the organizers and participants of the SpaceHack 3.0 hackathon, especially participants of this project: Alexander Sudy, Amos Münch, Jieran Sun, Mehdi Joodak, Kai Li, Samaneh Samiei, Darius Schaub, Siao-Han Wong, Tengyu Zhang.

## References

1. Long, Y. et al. Deciphering spatial domains from spatial multi-omics with SpatialGlue. Nat. Methods 21, 1658–1667 (2024).

2. Yuan, Z. et al. Benchmarking spatial clustering methods with spatially resolved transcriptomics data. Nat. Methods 21, 712–722 (2024).

3. Müller-Bötticher, N., Sahay, S., Eils, R. & Ishaque, N. SpatialLeiden: spatially aware Leiden clustering. Genome Biol. 26, 24 (2025).

4. Luecken, M. D. et al. Benchmarking atlas-level data integration in single-cell genomics. Nat. Methods 19, 41–50 (2022).

5. Long, Y. et al. Spatially informed clustering, integration, and deconvolution of spatial transcriptomics with GraphST. Nat. Commun. 14, 1155 (2023).

6. Vandereyken, K., Sifrim, A., Thienpont, B. & Voet, T. Methods and applications for single-cell and spatial multi-omics. Nat. Rev. Genet. 24, 494–515 (2023).

7. Merritt, C. R. et al. Multiplex digital spatial profiling of proteins and RNA in fixed tissue. Nat. Biotechnol. 38, 586–599 (2020).

8. Su, J.-H., Zheng, P., Kinrot, S. S., Bintu, B. & Zhuang, X. Genome-Scale Imaging of the 3D Organization and Transcriptional Activity of Chromatin. Cell 182, 1641-1659.e26 (2020).

9. Takei, Y. et al. Integrated spatial genomics reveals global architecture of single nuclei. Nature 590, 344–350 (2021).

10. Vickovic, S. et al. SM-Omics is an automated platform for high-throughput spatial multiomics. Nat. Commun. 13, 795 (2022).

11. Zhang, D. et al. Spatial epigenome–transcriptome co-profiling of mammalian tissues. Nature 616, 113–122 (2023).

12. Ben-Chetrit, N. et al. Integration of whole transcriptome spatial profiling with protein markers. Nat. Biotechnol. 41, 788–793 (2023).

13. Liu, Y. et al. High-plex protein and whole transcriptome co-mapping at cellular resolution with spatial CITE-seq. Nat. Biotechnol. 41, 1405–1409 (2023).

14. Zeng, H. et al. Integrative in situ mapping of single-cell transcriptional states and tissue histopathology in a mouse model of Alzheimer’s disease. Nat. Neurosci. 26, 430–446 (2023).

15. Liao, S. et al. Integrated Spatial Transcriptomic and Proteomic Analysis of Fresh Frozen Tissue Based on Stereo-seq. 2023.04.28.538364 Preprint at 10.1101/2023.04.28.538364 (2023).

16. Vicari, M. et al. Spatial multimodal analysis of transcriptomes and metabolomes in tissues. Nat. Biotechnol. 42, 1046–1050 (2024).

17. Guo, P. et al. Multiplexed spatial mapping of chromatin features, transcriptome and proteins in tissues. Nat. Methods 22, 520–529 (2025).

18. Duan, Z., Li, X., Xiao, Z., Ying, R. & Zhang, J. MUSE: A Multi-slice Joint Analysis Method for Spatial Transcriptomics Experiments. in Proceedings of the 34th ACM International Conference on Information and Knowledge Management 625–634 (Association for Computing Machinery, New York, NY, USA, 2025). doi:10.1145/3746252.3761240.

19. Lopez, R., Regier, J., Cole, M. B., Jordan, M. I. & Yosef, N. Deep generative modeling for single-cell transcriptomics. Nat. Methods 15, 1053–1058 (2018).

20. Korsunsky, I. et al. Fast, sensitive and accurate integration of single-cell data with Harmony. Nat. Methods 16, 1289–1296 (2019).

21. Polański, K. et al. BBKNN: fast batch alignment of single cell transcriptomes. Bioinformatics 36, 964–965 (2020).

22. Yao, Z. et al. A high-resolution transcriptomic and spatial atlas of cell types in the whole mouse brain. Nature 624, 317–332 (2023).

23. Wang, Q. et al. The Allen Mouse Brain Common Coordinate Framework: A 3D Reference Atlas. Cell 181, 936-953.e20 (2020).

24. Schott, M. et al. Open-ST: High-resolution spatial transcriptomics in 3D. Cell 187, 3953-3972.e26 (2024).

25. Bredikhin, D., Kats, I. & Stegle, O. MUON: multimodal omics analysis framework. Genome Biol. 23, 42 (2022).

26. Zheng, Y., Abila, E., Chrenková, E., Winkler, J. & Rendeiro, A. F. LazySlide: accessible and interoperable whole slide image analysis. 2025.05.28.656548 Preprint at 10.1101/2025.05.28.656548 (2025).

27. Oliveira, M. F. de et al. High-definition spatial transcriptomic profiling of immune cell populations in colorectal cancer. Nat. Genet. 57, 1512–1523 (2025).

28. Sun, J. et al. Beyond benchmarking: an expert-guided consensus approach to spatially aware clustering. 2025.06.23.660861 Preprint at 10.1101/2025.06.23.660861 (2025).

29. Townes, F. W. & Engelhardt, B. E. Nonnegative spatial factorization applied to spatial genomics. Nat. Methods 20, 229–238 (2023).

30. Virshup, I., Rybakov, S., Theis, F. J., Angerer, P. & Wolf, F. A. anndata: Access and store annotated data matrices. J. Open Source Softw. 9, 4371 (2024).

31. Wolf, F. A., Angerer, P. & Theis, F. J. SCANPY: large-scale single-cell gene expression data analysis. Genome Biol 19, (2018).

32. Palla, G. et al. Squidpy: a scalable framework for spatial omics analysis. Nat. Methods 19, 171–178 (2022).

33. Zhang, M. et al. Molecularly defined and spatially resolved cell atlas of the whole mouse brain. Nature 624, 343–354 (2023).

34. Chen, R. J. et al. Towards a general-purpose foundation model for computational pathology. Nat. Med. 30, 850–862 (2024).

35. Pedregosa, F. et al. Scikit-learn: Machine Learning in Python. J. Mach. Learn. Res. 12, 2825–2830 (2011).

36. Maynard, K. R. et al. Transcriptome-scale spatial gene expression in the human dorsolateral prefrontal cortex. Nat. Neurosci. 24, 425–436 (2021).

