## Supplementary Information for "Atlas-scale spatially aware clustering with support for 3D and multimodal data using SpatialLeiden"

### Supplementary Methods

#### Simulated multimodal data

##### SpatialLeiden

For SpatialLeiden the simulated data was preprocessed as described in the methods. The data was zero-centred, and 30 and 10 principal components (PCs) used for the RNA and protein modalities, respectively. Clustering was performed with spatialleiden_multimodal with a resolution of 0.5 and the layer weights set to 1 for protein, 1.5 for RNA, and 2.5 for the spatial layer.

##### SpatialGlue

SpatialGlue (spatialglue v1.1.5 with Python v3.8.20, scikit-learn v1.1.3, torch v2.4.1, and mclust v5.4.1) was applied to the simulated dataset following the tutorial. Briefly, for the RNA data 3,000 highly variable genes were identified (scanpy.pp.highly_variable_genes with flavor 'seurat_v3'), each cell normalized to $10^4$ counts (scanpy.pp.normalize_total), the counts log-transformed (scanpy.pp.log1p), and PCA calculated (SpatialGlue.preprocess.pca) for the first 100 PCs (corresponding to the number of proteins). The protein data was processed normalized (SpatialGlue.preprocess.clr_normalize_each_cell) and also PCA transformed using all PCs (SpatialGlue.preprocess.pca). Spatial neighbours were identified (SpatialGlue.preprocess.construct_neighbor_graph) and the SpatialGlue model trained for 200 epochs. Spatial domains were defined by clustering with mclust.

### Supplementary Figures

#### Supplementary Figure 1

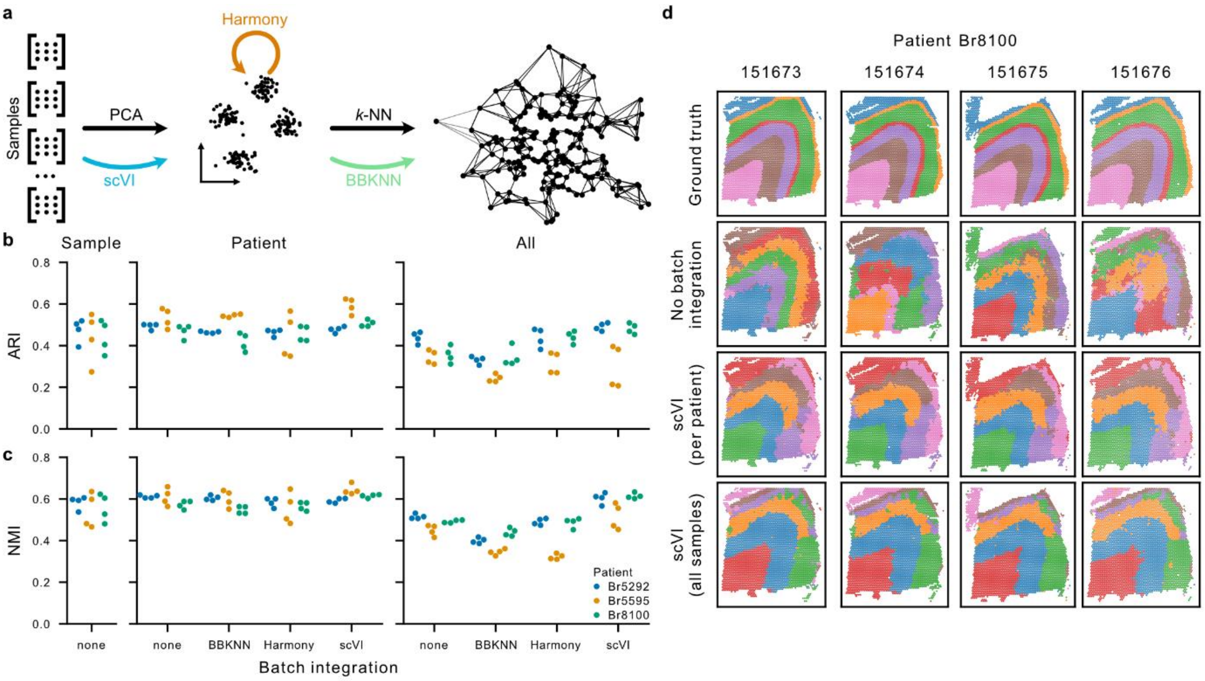

**Supplementary Figure 1: Batch integration enables multi-sample domain identification using SpatialLeiden. a,** Schematic indicating the different steps in a generalized workflow where batch integration methods can be applied. **b,** Adjusted Rand index (ARI) and **c,** normalized mutual information (NMI) when integrating and clustering the samples separately (left), patient-wise (middle), and across patients (right) in the DLPFC dataset. **d,** Ground truth and identified domains for patient Br8100 without integration and for scVI-integrated samples.

Supplementary Figure 2

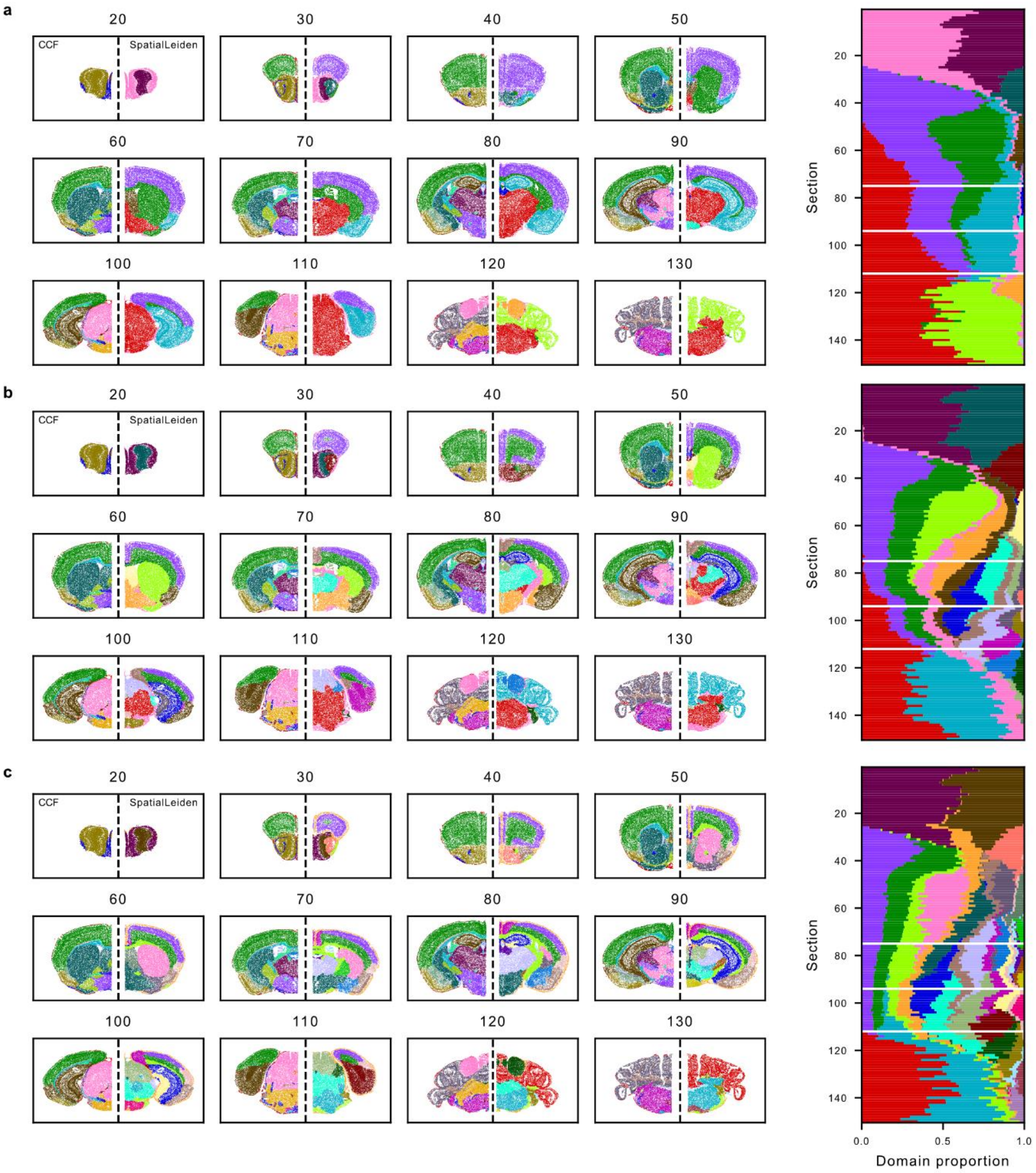

**Supplementary Figure 2: Increasing cluster number enables refinement of multi-scale spatial structures. a-c,** Example sections highlighting domains identified with SpatialLeiden (left) and proportion of cells per section (right) using a resolution of 0.5 (**a**), resolution of 1 (**b**), and resolution of 1.5 (**c**).

Supplementary Figure 3

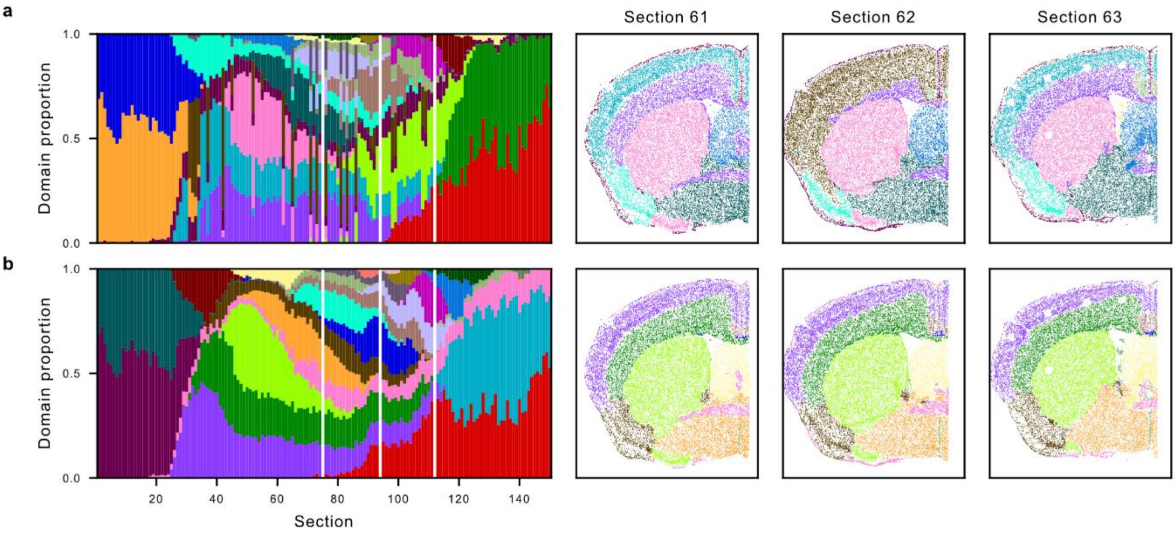

**Supplementary Figure 3: Batch integration improves spatial continuity of domains. a–b,** Proportions of cells per section belonging to each domain (left) and three consecutive slices (right) for SpatialLeiden without batch integration (**a**) and batch integration with scVI (**b**) highlight improved continuity of domains across sections after batch integration.

### Supplementary Figure 4

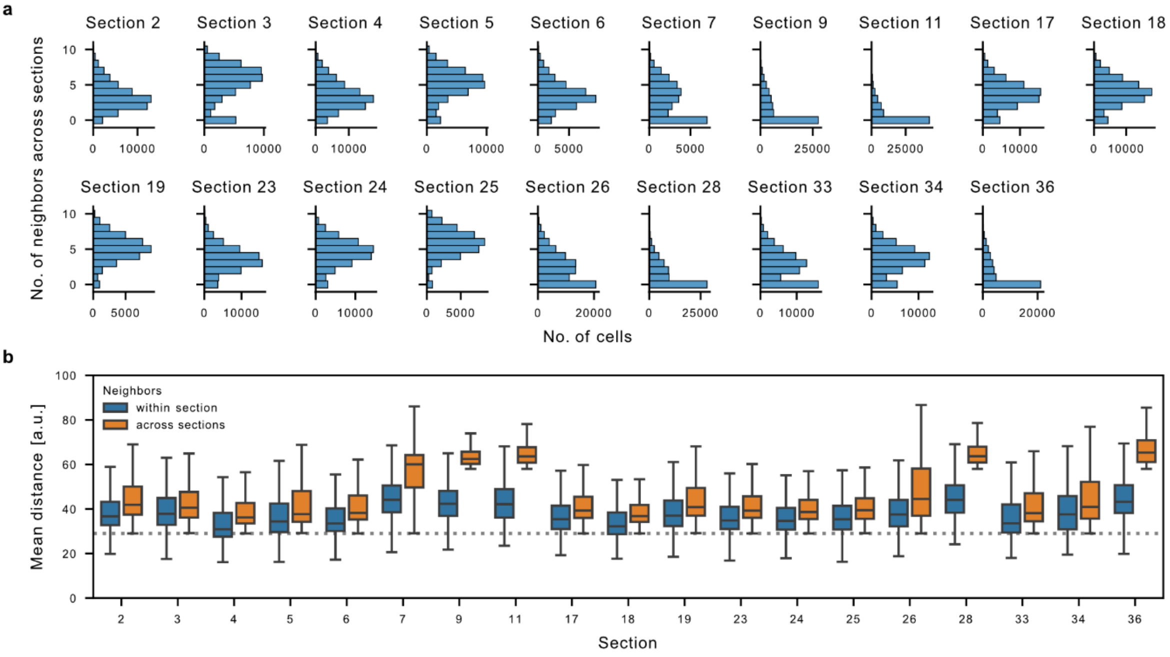

**Supplementary Figure 4: Spatial neighbours in Open-ST lymph node data stretch across adjacent sections. a,** Most cells have spatial neighbours in adjacent sections. **b,** Mean distance of each cell's neighbours within the same section and across sections. Dotted line indicates inter-section distance. Box plots show the median and the 25$^{th}$–75$^{th}$ percentiles, with whiskers extending to the most extreme data points within 1.5× the interquartile range.

### Supplementary Figure 5

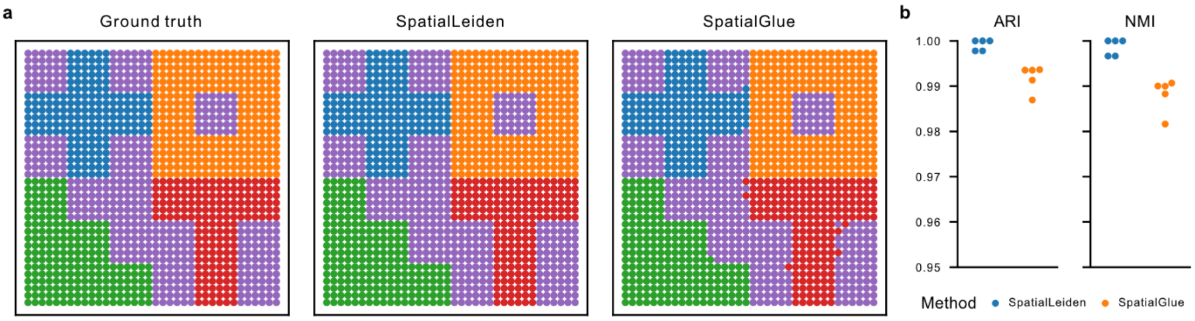

**Supplementary Figure 5: SpatialLeiden integrates simulated multimodal data. a,** Ground truth and domains identified by SpatialLeiden and SpatialGlue based on simulated dataset 1. **b,** Adjusted Rand index (ARI) and normalized mutual information (NMI) for five simulation datasets.

### Supplementary Figure 6

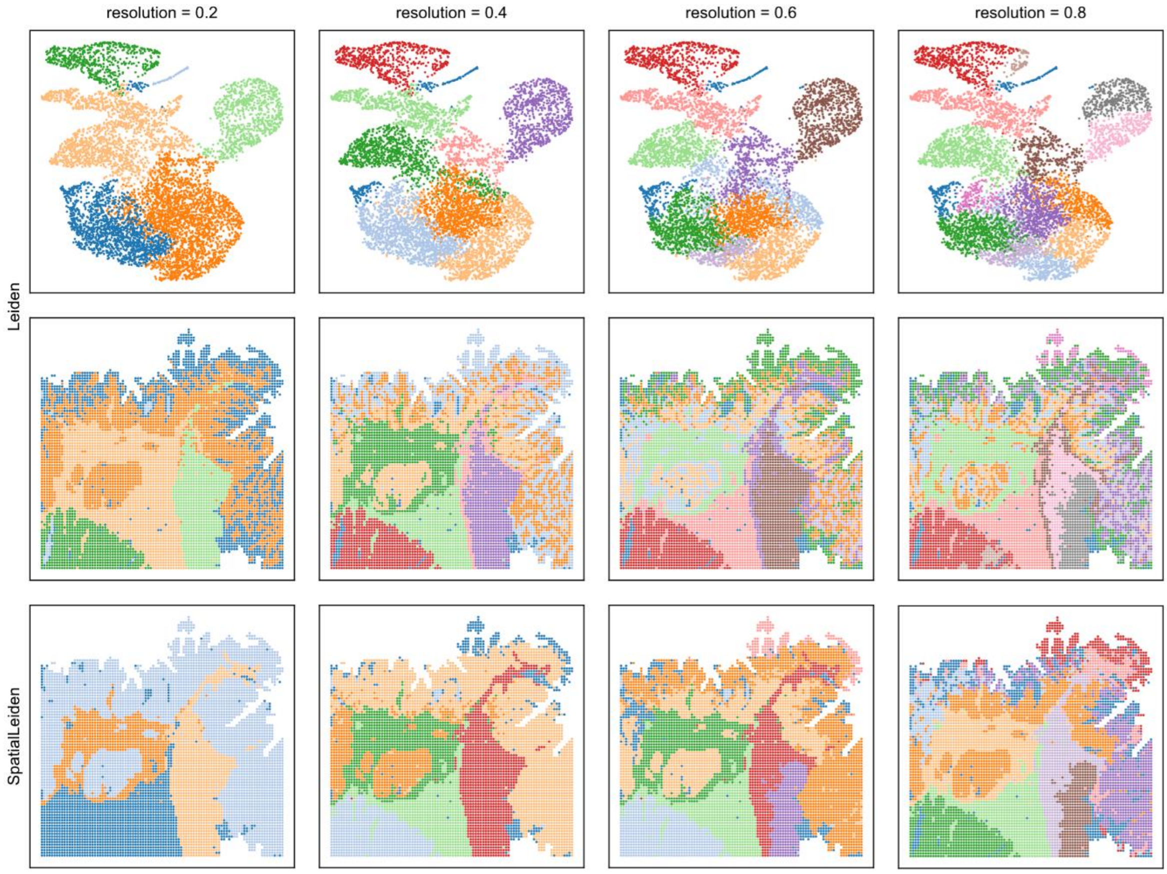

**Supplementary Figure 6: Leiden and SpatialLeiden clustering on H&E features using multiple resolutions.** Spatially aware and non-spatial Leiden clustering identifies smooth muscle, epithelial, and neoplastic compartments from H&E-stained slides.

Supplementary Figure 7

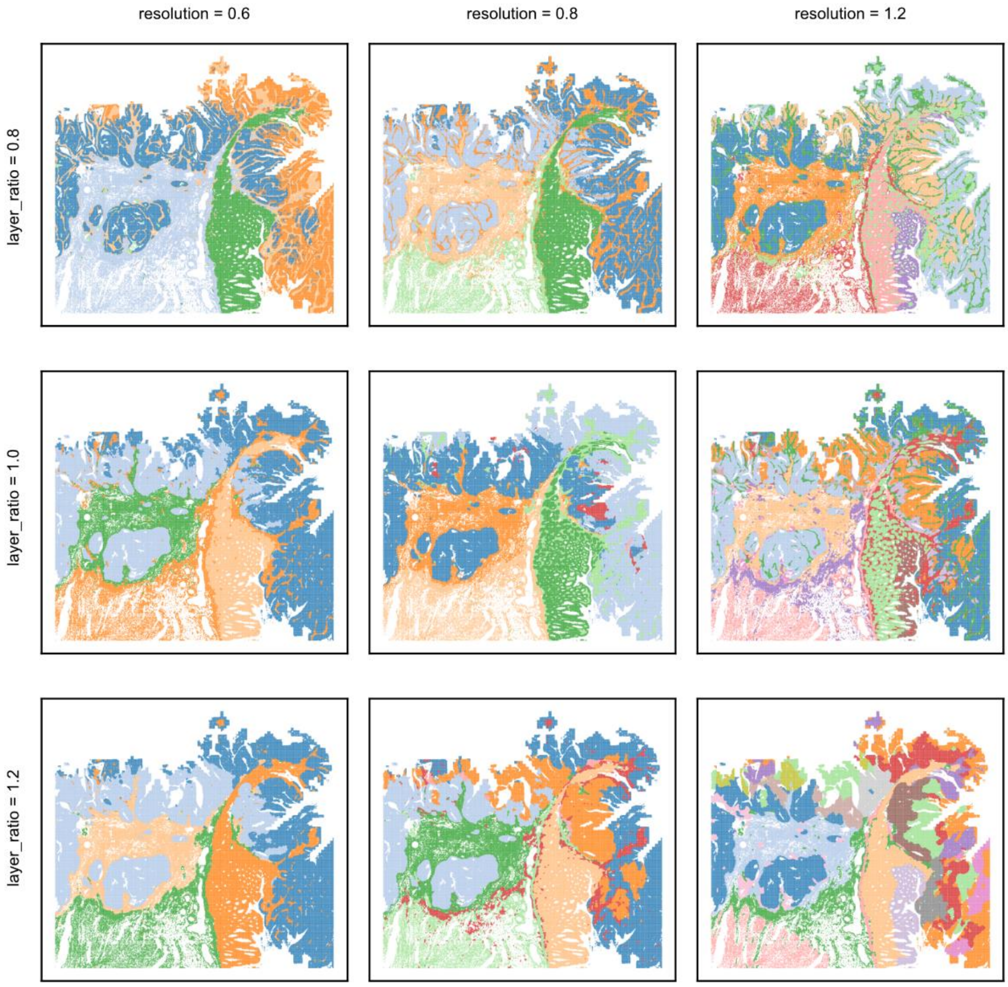

**Supplementary Figure 7: SpatialLeiden clustering on Visium HD data using multiple resolutions and layer ratios.** SpatialLeiden clustering identifies epithelial, neoplastic, and dysplastic compartments from spatially resolved transcriptomics data, whereas the smooth muscle compartment is identifiable only through over clustering.
